# Apolipoprotein E4 modulates astrocyte neuronal support functions in the presence of amyloid-β

**DOI:** 10.1101/2022.09.29.510145

**Authors:** Rebecca M Fleeman, Madison K Kuhn, Dennis C Chan, Elizabeth A Proctor

## Abstract

Apolipoprotein E (APOE) is a lipid transporter produced predominantly by astrocytes in the brain. The ε4 variant of APOE (APOE4) is the strongest and most common genetic risk factor for Alzheimer’s disease (AD). Although the molecular mechanisms of this increased risk are unclear, APOE4 is known to alter immune signaling and lipid and glucose metabolism. Astrocytes provide various forms of support to neurons, including regulating neuron metabolism and immune responses through cytokine signaling. Changes in astrocyte function due to APOE4 may therefore decrease neuronal support, leaving neurons more vulnerable to stress and disease insults. To determine whether APOE4 alters astrocyte neuronal support functions, we measured glycolytic and oxidative metabolism of neurons treated with conditioned media from APOE4 or APOE3 (the common, risk-neutral variant) primary astrocyte cultures. We found that APOE4 neurons treated with conditioned media from resting APOE4 astrocytes had similar metabolism to APOE3 astrocytes, but treatment with ACM from astrocytes challenged with amyloid-β (Aβ), a key pathological protein in AD, caused APOE4 neurons to increase their basal mitochondrial and glycolytic metabolic rates more than APOE3 neurons. These changes were not due to differences in astrocytic lactate production or glucose utilization, but instead correlated with increased glycolytic ATP production and a lack of cytokine secretion response to Aβ. Together, these findings suggest that in the presence of Aβ, APOE4 astrocytes alter immune and metabolic functions that result in a compensatory increase in neuronal metabolic stress.

## 1. Introduction

Apolipoprotein E (APOE) is a lipid transporter found in the brain and the periphery, and is important for shuttling lipid molecules between cells to maintain membranes, modulate growth of cellular projections, and repair injured cells^1,2^. Humans express three common APOE isoforms, APOE ε2, ε3, and ε4, which differ in only two amino acid residues^3,4^. The majority (78% of humans) express APOE3, while APOE4 is the greatest and most common genetic risk factor for Alzheimer’s disease (AD)^5–8^. Although only 15% of people are APOE4 carriers, 60% or more of AD patients carry at least one copy of APOE4^5,9^. While the pleiotropic effects of APOE4 have been examined in multiple cell types, the mechanism by which these effects may increase AD risk is not fully understood^1,6,10^.

Astrocytes are glial cells that support neural homeostasis through regulating neuron metabolism, maintaining synaptic connectivity, supplying neurotransmitters, and modulating immune response via cytokine signaling^11–14^. Astrocytes are the primary producers of APOE in the brain^15^. APOE shuttles cholesterol to and from neurons for maintenance of axonal growth, synaptic formation, and transport, as well as neurotransmitter release^1,15–17^. APOE isoform directly affects the astrocytes that produce it, including APOE genotype-dependent alterations in astrocytic lipid metabolism^18^, APOE secretion^19^, and immune responses^20^. These differences remain incompletely understood, and how these APOE-isoform-specific changes in metabolism and reactivity of astrocytes may negatively affect neuron health and promote disease is an open question in the field.

AD is a neurodegenerative disease characterized by two proteinopathies: the deposition of amyloid-β (Aβ) plaques and neurofibrillary tau tangles^21^. While many healthy aging brains will accumulate Aβ and tau inclusions, the correlation between plaque/tangle load and neurodegeneration or cognitive impairment is only modest^22^. We hypothesize that APOE4-associated lack of astrocytic support creates vulnerabilities in neurons, leaving neurons more susceptible to stress-induced death when challenged with AD proteinopathies. Identifying the mechanisms by which APOE4 reduces astrocytic support of neurons will open new avenues of investigation in therapeutically mitigating APOE4-associated AD risk.

Here, we focus on how APOE4 affects astrocytic and neuronal metabolism, and how Aβ stimulation of astrocytes affects the immune and metabolic support astrocytes provide neurons. We find that APOE genotype determines the response of astrocytes to Aβ, where APOE4 astrocytes lack an Aβ-stimulus cytokine signaling response but upregulate glycolytic ATP production. In turn, this Aβ-stimulated APOE4 astrocytic response increased APOE4 neuronal mitochondrial and glycolytic metabolism. These findings indicate that APOE4 creates subtle shifts in reactive astrocytic responses to Aβ, which stresses neuronal metabolism, resulting in potential susceptibility to disease insults.

## 2. Methods

### 2.1 Primary neuron and astrocyte culture

Animal protocols were approved by the Penn State College of Medicine Institutional Animal Care and Use Committee (PROTO201800531). The mice used in this study are humanized APOE knock-in animals carrying the APOE3 (B6.Cg-Apoe^em2(APOE*)Adiuj^/J) or APOE4 (B6(SJL)-Apoe^tm1.1(APOE*4)Adiuj^/J) variants (Jackson Laboratory, Bar Harbor, ME, USA). APOE4 animals were purchased as homozygotes, while APOE3 animals were purchased as heterozygotes and bred to homozygosity, with homozygosity confirmed by qPCR. Breeders were fed a standard chow diet *ad libitum* (Teklad 2018, Envigo). We generated primary cell cultures from postnatal day 0 or day 1 (P0/1) APOE3 and APOE4 pups. Each biological replicate was an individual pup from 3 or more separate litters. Each pup brain was kept separate and given its own plate. P0/1 pups were sacrificed by decapitation using surgical scissors. Isolated brains were placed in cold HEPES-buffered Hanks’ Balanced Salt solution (pH 7.8). The brain was dissected to isolate the cortical cap of each hemisphere, and meninges were removed.

For neuron culture, isolated cortices were transferred to conical tubes of warm embryonic neuron plating medium: Neurobasal Plus (Gibco), 10% FBS (Gibco), 1× GlutaMAX (Gibco), 1× Penicillin-Streptomycin (10,000 U/mL, Gibco). Cortices were triturated in plating medium with a p1000 pipette. Cell concentration was measured using the Countess II automated cell counter (Invitrogen) and cells were plated at 154,500 cells/well on poly-D-lysine (Gibco)-coated 24-well Seahorse plates. The cells kept at 37°C, 5% CO_2_ to allow cells to attach. 2-12 hours after plating, medium was switched to Neuronal Medium: Neurobasal Plus (Gibco), 1X B27 Plus supplement (Gibco), 1X GlutaMAX (Gibco), 1X Penicillin-Streptomycin (10,000 U/mL, Gibco). Half the medium was changed every 5-7 days. Neurons were treated with astrocytic conditioned media on day 11 and assayed in Seahorse XFe24 on day 14.

For astrocyte culture, isolated cortices were transferred to conical tubes of warm glial plating medium: DMEM/F12, 20% FBS, 5 mL Penn/Strep, 5 mL 100mM Sodium Pyruvate, 50 μL 0.1 mg/mL EGF stock in ddH_2_O. Cortices were triturated in plating medium with a p1000 pipette then transferred to a T-25 cell culture flask coated with Poly-D-Lysine (Gibco) with additional glial media for a total volume of 4 mL. Media was replaced every 2-3 days. Once confluent, the flask shook at 180 rpm for 2 hours and media containing non-adherent glial cells was removed. Astrocytes were then washed with 1X phosphate buffered saline (PBS), removed with trypsin-EDTA, quenched with glial media, and plated in a Poly-D-Lysine coated 6-well or Seahorse plate at 1,000,000 and 50,000 cells per well, respectively. Once 75% confluent, cells were treated with corresponding conditions.

### 2.2 Amyloid-β Aggregation & Addition

One milligram of human Aβ_42_ (Novex 75492034A) was solubilized in 100 μL Trifluoroacetic acid (TFA) and aggregated by diluting to 22.15 μM with PBS, followed by incubation to induce aggregation at 37° C for 24 hours. We added Aβ aggregates to astrocytes diluted in fresh glial media at 1 μM. After 72 hours, media was removed, flash frozen in liquid nitrogen, and stored at −80° C until use.

### 2.3 Metabolic Analysis

Mitochondrial and glycolytic function was analyzed with the Seahorse Cell Mito Stress Test and Seahorse Real-Time ATP Rate Assay on a Seahorse XFe24 Extracellular Flux Analyzer (Agilent). Neurons were treated on day 11 of culture with 500 μL of fresh neuronal media plus 500 μL of astrocytic conditioned media of varying treatment. This 50/50 mix of fresh neuronal media and ACM treatment ensured enough nutrients were provided to sustain neuronal viability. Astrocytes were treated once 75% confluent with 1 mL of media containing either 1 μM Aβ or equivalent amount of vehicle, or fresh media controls. For both cell types, 72 hours after treatment, media was removed and flash frozen in liquid nitrogen. Media was replaced with 500 μL of seahorse assay media (Seahorse XF DMEM or RPMI medium, 1 mM pyruvate, 2 mM glutamine, and 10 mM glucose, pH 7.4). Oxygen consumption rate (OCR) was measured at baseline and after addition of 1.5 μM oligomycin (Complex V inhibitor), 1 μM FCCP (uncoupler), and 0.5 μM rotenone/antimycin A (Complex I and Complex III inhibitors, respectively) for the Mito Stress Test, and 1.5 μM oligomycin and 0.5 μM rotenone/antimycin A for the ATP Rate Assay. Final OCR was normalized by protein concentration (Pierce™ BCA Protein Assay (Thermo Scientific)) and to non-mitochondrial respiration to account for plate-to-plate variation and batch effects^23^. All neuron Seahorse results are averages of six or more biological replicates (different pups from multiple litters) and each biological replicate consisted of the median of three to six technical replicates (wells per plate). All astrocyte Seahorse results are averages of three or more biological replicates and each biological replicate consisted of the median of three to five technical replicates.

### 2.4 Detection of Glucose and Lactate Concentration

Glucose levels in cell culture media were measured by GlucCell Glucose Monitoring System (CESCO Bioengineering) following the manufacturer’s protocol. The neuronal and glial media have glucose concentrations of 450 mg/dL and 315.1 mg/dL, respectively, and the meter’s range is 30-500 mg/dL. Lactate levels in cell culture media were measured by L-Lactate Assay Kit (Cayman Chemical) following the manufacturer’s protocol. The neuronal media does not contain lactate; the glial media does contain lactate in the mM range due to FBS. Thus, we diluted all samples 50× to maintain samples within the lactate assay’s sensitivity range of 25 μM to 1mM.

### 2.5 Cytokine Concentration

We quantified the levels of 32 cytokines on the Luminex FLEXMAP 3D platform using a MILLIPLEX Mouse Cytokine/Chemokine Magnetic Bead Panel (MCYTOMAG-70K) according to the manufacturer’s protocol, accommodating to a 384-well plate format, where magnetic beads and antibodies were used at half-volume. All samples were thawed on ice and assayed in technical triplicate, with n of 9-10 samples per group.

We used an in-house automated data cleaning pipeline, available on our laboratory’s GitHub page (https://github.com/elizabethproctor/Luminex-Data-Cleaning) to prepare the cytokine concentration data for analysis. Briefly, we first reviewed the bead counts of all observations, with an overall average bead count of 70 beads/well. Next, we calculated pairwise differences in fluorescence measurements within each technical triplicate. If the separation of the replicate was greater than twice the distance between the other two, it was designated an outlier and removed before further analysis. After excluding outliers (2% of measurements), we calculated the average of the remaining technical replicates for PLS analysis.

### 2.6 Partial Least Squares Discriminant Analysis

We mean-centered and unit-variance scaled (*i.e*., Z-scored) the cytokine data prior to performing partial least squares (PLS) analysis. PLS X-block (predictors) were cytokine concentrations and Y-block (outputs) were genotype or Aβ treatment. We performed PLS in R (v.3.6.1) using ropls (v.1.16.0)^24^ and plotted with ggplot2 (v.3.2.1)^25^. We calculated accuracy by performing cross-validation with one-third of the data and calculated model confidence by comparing the predictive accuracy of cross-validation from our model to the distribution of crossvalidation accuracies of 100 randomized models. We constructed these randomized models by randomly permuting the class assignment to the preserved X-block, conserving the data landscape to provide a valid control.

We orthogonalized each PLS model on the first latent variable (LV1), so that the parameter variance that most correlated with our chosen grouping was maximally projected onto LV1. The number of latent variables for each model was chosen based on the model with the lowest cross-validation error, and this latent variable number was maintained while constructing the distribution of randomized models for confidence calculations. Variable importance in projection (VIP) scores were calculated by quantifying the contribution of each cytokine to the prediction accuracy of our model. We calculated VIP scores by averaging the weight of each cytokine on each latent variable in the model, normalized by percent variance explained by each respective latent variable:

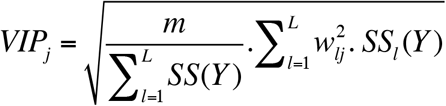

where *m* is the total number of predictors, *l* is the latent variable, *L* is the number of latent variables in the model, *w_lj_* is the weight (inverse loading) of predictor *j* on latent variable *l*, and *SS_l_*(*Y*) is the variation in class Y explained by latent variable *l*. Due to normalization, the average of squared VIP scores is 1, meaning that a VIP > 1 designates a parameter as having an above average contribution to our model.

### 2.7 Statistical Analysis

All data are expressed as mean ± standard error of the mean. Statistical analyses were conducted in Graph Pad Prism 9 (v 9.4.1). T-tests were used to compare basal neuron OCR, ECAR, and Aβ concentration between genotypes. We used 2-way ANOVA to determine differences between astrocyte glucose and lactate in 3 conditions. We tested for outliers using the ROUT method with Q of 0.5. Statistical significance was determined using an error probability level of p < 0.05.

## 3. Results

### 3.1 Aβ-stressed APOE4 astrocytes promote increased neuronal metabolism

APOE genotype directly affects lipid and glucose metabolism^18,26,27^. To first confirm the effects of APOE4 on neuronal mitochondrial and glycolytic metabolism, we cultured APOE3 and APOE4 primary murine neurons to maturity and quantified mitochondrial function and glycolytic rate at rest. We found no differences in the mitochondrial respiration (oxygen consumption rate, OCR) (**Figure 1A**) or non-mitochondrial glycolysis (extracellular acidification rate, ECAR) of neurons (**Figure 1E**) due to APOE genotype, suggesting that APOE4 does not alter neuronal resting mitochondrial respiration.

**Figure 1.**
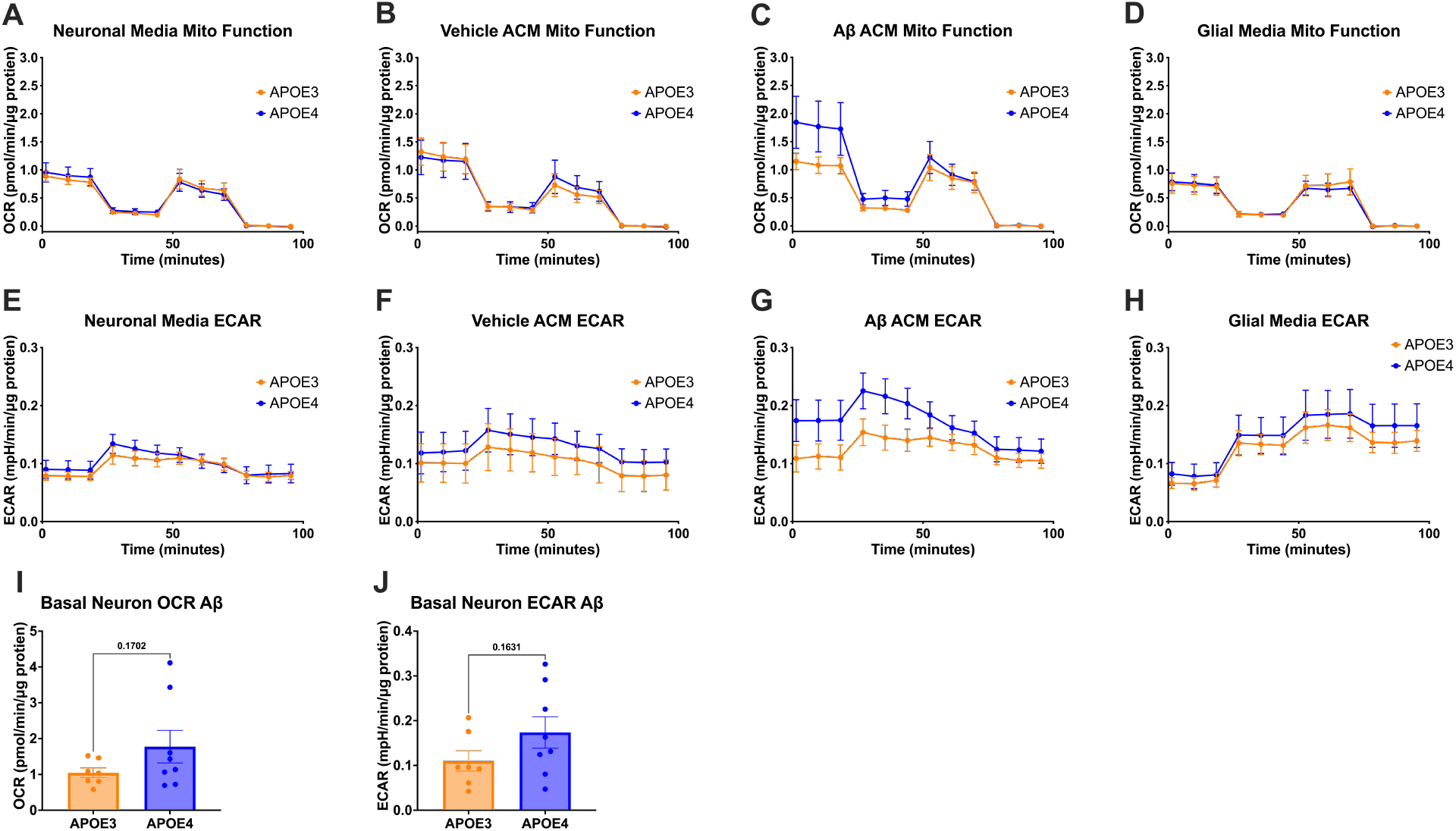
APOE4 neurons increase oxidative and glycolytic output in response to ACM from Aβ-treated astrocytes. Mitochondrial stress test oxygen consumption rates (OCR) of APOE3 and APOE4 neurons treated for 72 hours with (A) fresh neuronal media, (B) Vehicle-treated astrocyte conditioned media (ACM), (C) 1 μM Aβ-treated ACM, or (D) glial media that had not been on cells. Neurons were treated with sequential injections of mitochondrial inhibitors oligomycin A, FCCP, and rotenone/antimycin A. Corresponding extracellular acidification rates (ECAR) of mitochondrial stress tests of APOE3 and APOE4 neurons treated for 72 hours with (E) fresh neuronal media, (F) Vehicle-treated ACM, (G) 1 μM Aβ-treated ACM, or (H) glial media that had not been on cells. Quantifications of basal OCR (I) and ECAR (J) of neuronal mitochondrial stress tests.

Because astrocytes are the primary producers of APOE and are uniquely situated to impact neuronal metabolism due to their support of immune, metabolic, synaptic, and trophic function, we next measured how treating neurons with astrocytic conditioned media (ACM) would affect neuronal metabolism. We cultured APOE3 and APOE4 primary murine astrocytes to maturity and treated neurons with their corresponding genotype ACM for 72 hours. We again saw no change in mitochondrial respiration due to APOE genotype in the neurons (**Figure 1B and 1F**).

With APOE4 being the greatest genetic risk factor for AD, we next asked how stimulating astrocytes with Aβ, the primary pathological protein in AD, may cause genotype-specific changes in the factors they secrete, downstream impacting neuronal metabolism. Aβ is known to have both direct and indirect neurotoxic effects, and Aβ-specific alterations in astrocytic function have the potential to negatively impact neuronal health. We treated mature primary APOE3 and APOE4 astrocytes with 1 μM Aβ for 72 hours and collected the ACM. Then we treated mature APOE3 and APOE4 primary neurons with media from Aβ-treated astrocytes (**Figure 1C and 1G**), or with glial media without cells, to rule out effects of the factors present in the glial media (**Figure 1D and 1H**). APOE4 neurons exhibited 70% higher basal mitochondrial respiration than APOE3 Aβ ACM on APOE3 neurons (**Figure 1C and 1I**), as well as a 58% higher non-mitochondrial glycolytic rates (**Figure 1G and 1J**), in response to the astrocytic Aβ stimulation secreted factors.

### 3.2 APOE genotype does not affect astrocytic lactate production or glucose utilization

To identify why APOE4 neurons treated with media from Aβ-treated astrocytes increase their basal metabolism, we first asked how much of the Aβ was left in the ACM after treatment, which would then be included in the neuron stimulation with ACM. After 72 hours of incubation with 1 μM Aβ, APOE3 and APOE4 astrocytes have similar levels of Aβ left in media (14 nM, or 64 ng/mL) (**Figure 2A**), indicating that both APOE3 and APOE4 astrocytes internalize 99% of the Aβ used for treatment. After 1:1 dilution with fresh neuronal media to ensure adequate levels of nutrients for neuron health, 7 nM of Aβ was present in the ACM added to neuron cultures, indicating that neurons were treated with only 0.7% of a neurotoxic dose of Aβ. Following the 72 hour treatment incubation, 3.5 nM of Aβ remained in the neuron culture media (**Figure 2B**). With no differences in Aβ levels observed in the media of APOE3 and APOE4 astrocytes or neurons, Aβ is an unlikely culprit for the observed differences in basal neuronal metabolism following exposure to ACM from Aβ-treated astrocytes.

**Figure 2.**
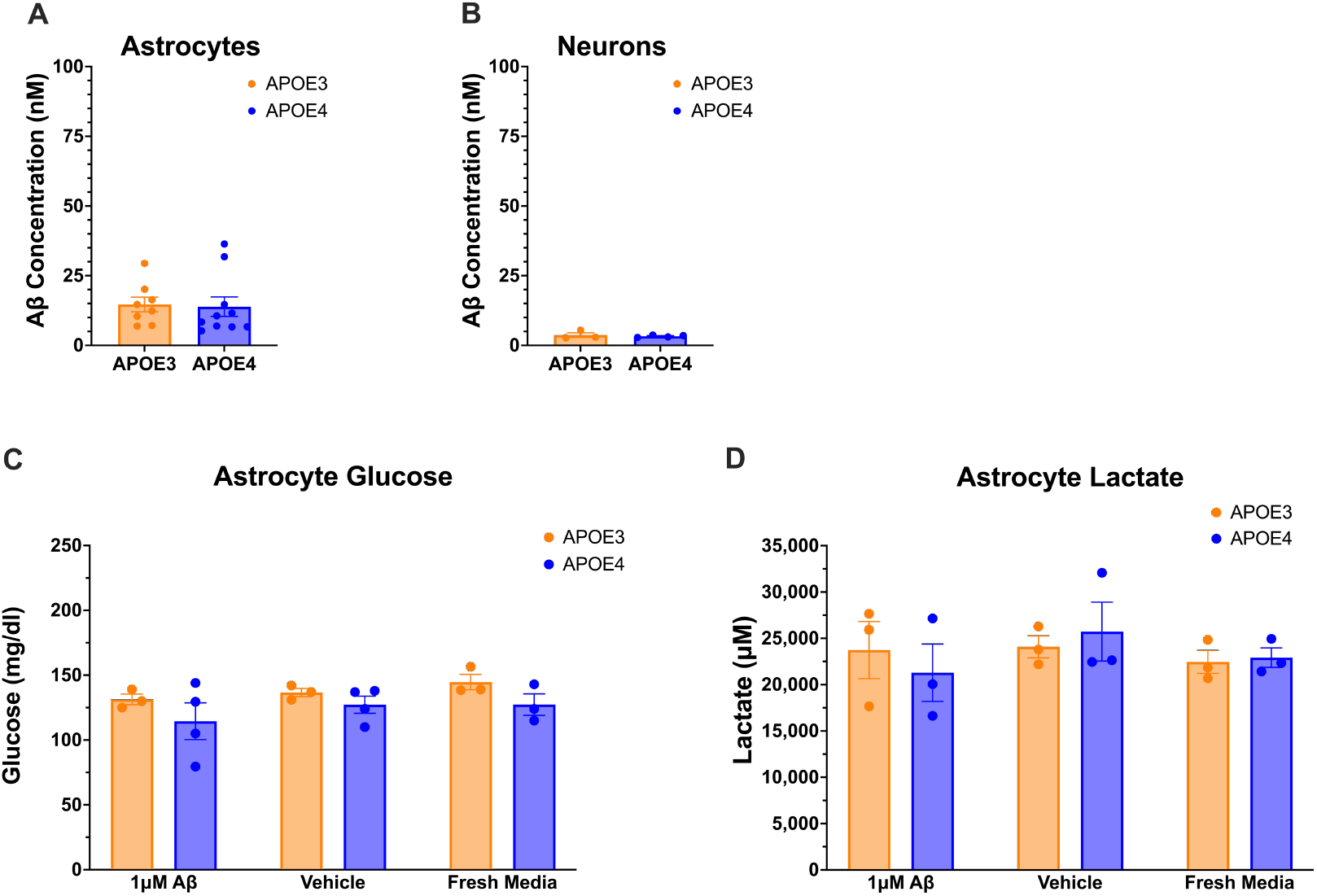
Aβ internalization, glucose consumption, and lactate production are similar in APOE3 and APOE4 astrocytes. (A) Concentration of Aβ remaining after 72-hour incubation of 1μM Aβ on astrocytes. (B) Concentration of Aβ remaining after 72-hour incubation of Aβ-treated ACM on neurons. Levels of glucose (C) and lactate (D) in astrocyte media after 72-hour treatment with 1 μM Aβ, Vehicle, or Fresh glial media.

We next asked whether Aβ-treated APOE4 astrocytes consume less glucose from the media than APOE3 astrocytes, resulting in higher levels of glucose in the ACM that might explain elevated basal metabolism in neurons. However, neither APOE genotype nor Aβ treatment appreciably affected glucose concentration in ACM (**Figure 2C**).

Finally, because astrocytes can provide lactate as a metabolic substrate to neurons, we asked whether elevated neuronal basal mitochondrial metabolism can be explained by higher lactate production by APOE4 astrocytes when treated with Aβ. However, similar to glucose, lactate levels were not altered by APOE genotype, nor by Aβ treatment (**Figure 2D**).

### 3.3 Aβ treatment increases APOE4 astrocyte glycolytic ATP production

We quantified the mitochondrial respiration and glycolytic rates of APOE3 and APOE4 astrocytes to better understand how differences in their metabolic function may influence neuron health. APOE4 astrocytes have lower basal and maximal respirations levels in comparison to APOE3 astrocytes, both at rest (**Figure 3B**) and in the presence of Aβ (**Figure 3A**), indicating that APOE4 astrocytes operate at a lower aerobic metabolic rate than APOE3 astrocytes. Additionally, Aβ treatment increased the maximal respiration, but not basal respiration, in both APOE3 and APOE4 astrocytes (**Figure 3C and 3D**). Similar to mitochondrial respiration, APOE4 astrocyte glycolytic output at rest was lower than that in resting APOE3 astrocytes (**Figure 3F**).

**Figure 3.**
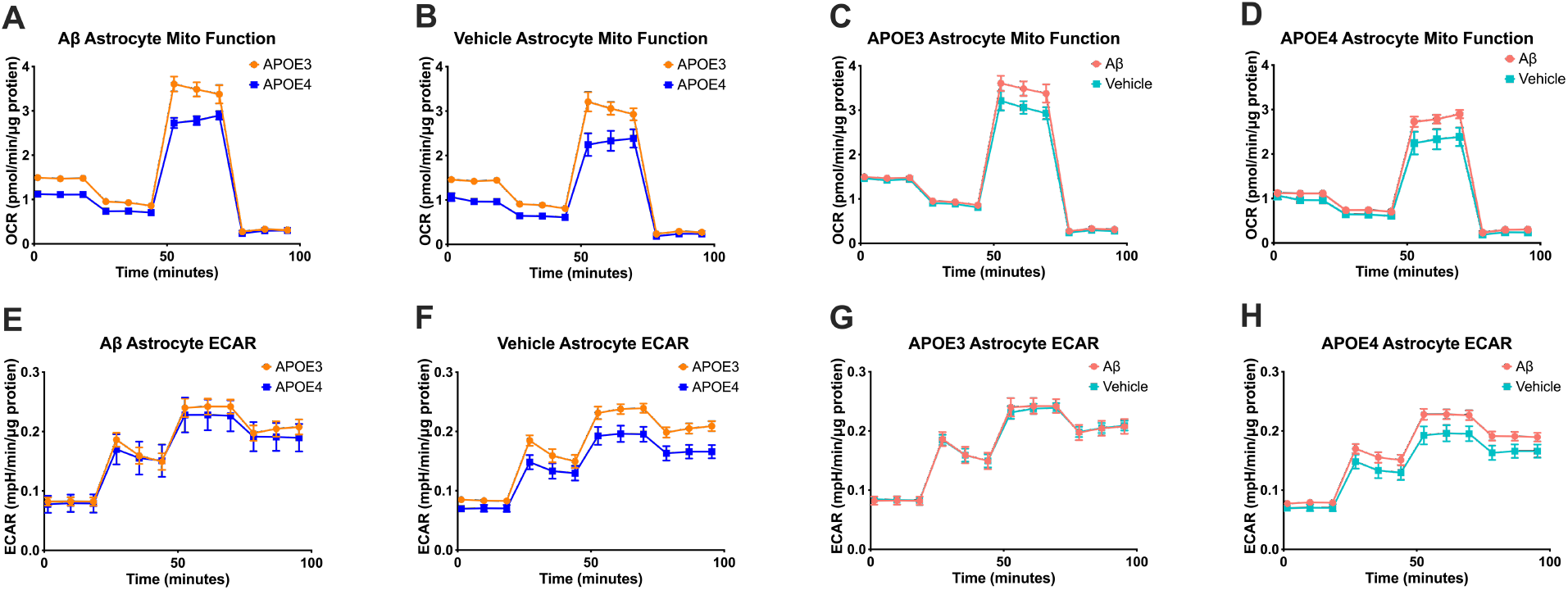
APOE4 astrocytes have lower metabolic function than APOE3 but have increased glycolysis response to Aβ. Mitochondrial stress test OCR (A and B) and ECAR (E and F) of APOE3 and APOE4 astrocytes treated for 72 hours with 1 μM Aβ or Vehicle. APOE3 mitochondrial stress test OCR (C) and ECAR (G) of astrocytes treated for 72 hours with 1 μM Aβ or Vehicle. APOE4 mitochondrial stress test OCR (D) and ECAR (H) of astrocytes treated for 72 hours with 1 μM Aβ or Vehicle.

However, the glycolytic rate in Aβ-treated APOE3 and APOE4 astrocytes was similar (**Figure 3E**). A comparison between treatments within genotypes shows that Aβ stimulation increased glycolysis in APOE4, but not APOE3, astrocytes (**Figure 3G and 3H**).

We next tested whether APOE4 astrocytes may be increasing APOE4 neuronal metabolism through differences in astrocytic ATP production in the presence of Aβ, since ATP is a key signaling molecule by which astrocytes modulate neuronal activity^28,29^. We quantified astrocyte ATP production rates from glycolysis and mitochondrial respiration and observed that APOE4 astrocytes produced nearly double the amount of glycolytic ATP as APOE3 astrocytes in the presence of Aβ, but lower levels at rest (**Figure 4A**). Hence, APOE4 astrocytes are more metabolically reactive to Aβ than are APOE3 astrocytes.

**Figure 4.**
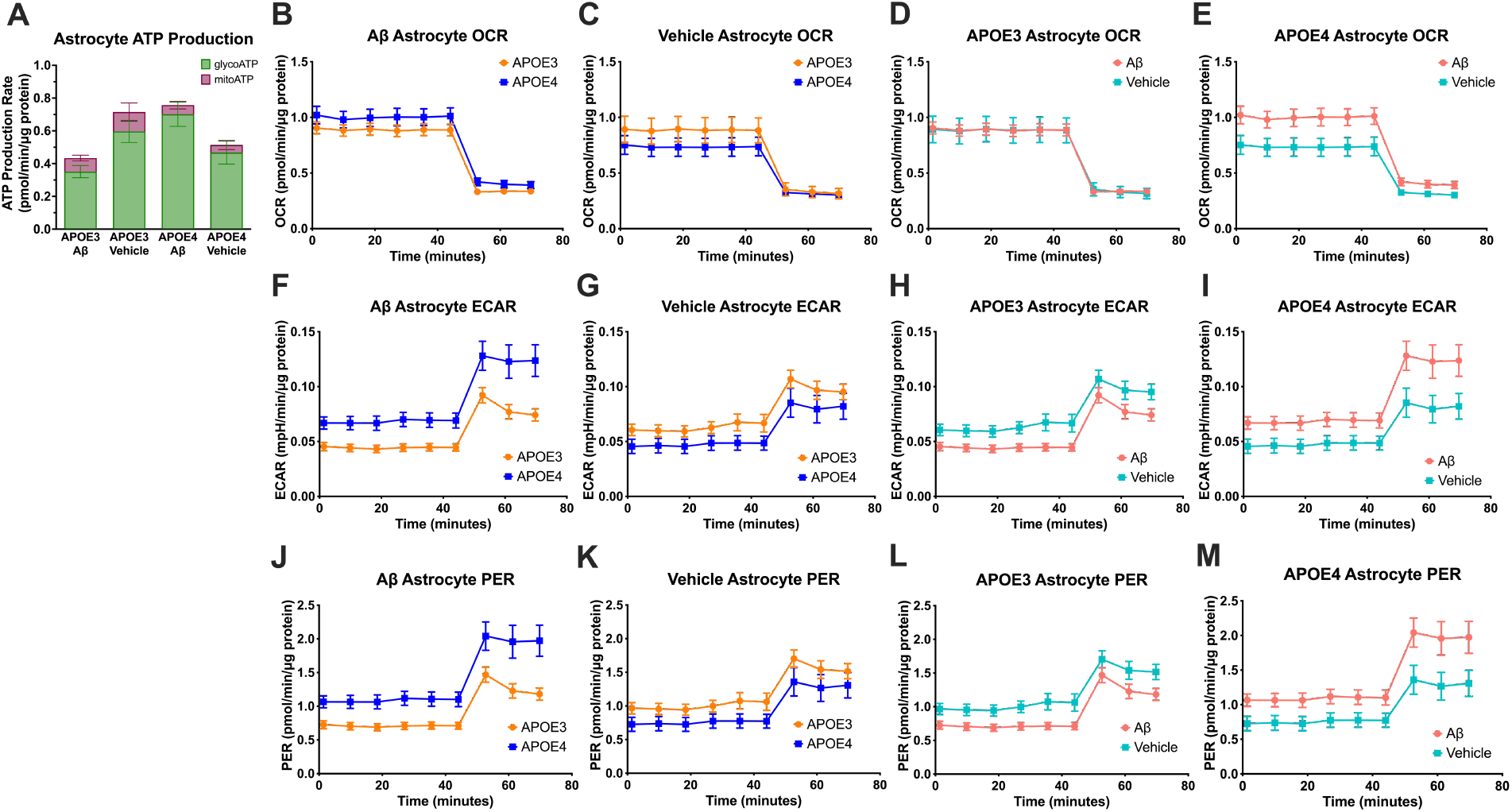
APOE4 astrocytes increase ATP production through glycolysis more than APOE3 astrocytes. (A) ATP production rate comparisons between APOE3 and APOE4 astrocytes treated with 1 μM Aβ or Vehicle. Seahorse ATP rate test on APOE3 and APOE4 astrocytes treated for 72 hours with 1 μM Aβ, OCR (B), ECAR (F), and proton efflux rate (PER) (J), with serial injections of mitochondrial inhibitors oligomycin A and rotenone/antimycin A. Seahorse ATP rate test on APOE3 and APOE4 astrocytes treated with Vehicle, OCR (C), ECAR (G), and PER (K). Seahorse ATP rate test on APOE3 astrocytes treated with 1μM Aβ or Vehicle, OCR (D), ECAR (H), and PER (K). Seahorse ATP rate test on APOE4 astrocytes treated with 1 μM Aβ or Vehicle, OCR (E), ECAR (I), and PER (L).

In addition to glycolytic ATP production, APOE4 astrocytes also increase mitochondrial glucose metabolism when exposed to Aβ (**Figure 4E**), while APOE3 astrocytes did not (**Figure 4D**). However, a direct comparison of glucose metabolism between APOE3 and APOE4 astrocytes showed no significant difference in mitochondrial glucose metabolism at rest nor in the presence of Aβ (**Figure 4B and 4C**).

We next quantified the extracellular acidification rates, which reflect production of free protons from ATP and lactate in glycolysis, and CO2 from mitochondrial respiration. We found that APOE4 astrocytes have higher acidification rates than APOE3 astrocytes in the presence of Aβ, but not at rest (**Figure 4F and 4G**). More specifically, Aβ decreased acidification in APOE3 astrocytes but increased acidification in APOE4 astrocytes (**Figure 4H and 4I**). Based on our finding of no changes in lactate production due to APOE genotype or Aβ treatment in astrocytes (**Figure 2D**), as well as the observation that ECAR is most prominently affected by Aβ after the addition of rotenone/antimycin A (Complex I and Complex III inhibitors, respectively, that shut down the electron transport chain of oxidative phosphorylation), our data strongly support our finding that APOE4 astrocytes increase glycolytic ATP production in the presence of Aβ. This increase is also reflected in an increase in proton efflux rate (PER) for APOE4 astrocytes treated with Aβ, but not APOE3 astrocytes (**Figures 4J-M**).

### 3.4 APOE4 astrocytes do not respond to Aβ stimulation with changes to cytokine secretion

In addition to the multitude of astrocytic metabolic pathways that impact neuronal health, cytokines secreted by glial cells also influence neuron metabolism^30^. APOE4 is thought to increase AD risk through several mechanisms, including by altering immune signaling networks^31^. Importantly, immune signaling involves many interacting pathways, necessitating multivariate analysis to properly account for interaction and covariation of cytokine pathways. Multivariate modeling uncovers small, highly correlated differences in cytokine levels that allow us to identify immune signaling “signatures.” Using the supervised machine learning tool partial least squares (PLS), we identified patterns of astrocytic cytokine secretion associated with each APOE genotype.

After quantifying the cytokine levels in conditioned media of mature primary APOE3 and APOE4 astrocytes using multiplexed immunoassays, we used PLS to predict APOE genotype based on resting levels of cytokine secretion (**Figure 5A and 5B**). APOE genotype has a clear effect on astrocyte immune signaling, even in the absence of AD pathology, as evidenced by the 70% cross-validation accuracy and 95% confidence in our ability to predict genotype using our model. APOE3 astrocytes secreted significantly higher levels of TNFα, MIP-1β, MCP-1, KC, IP-10, IL-4, and IL-5, while APOE4 astrocytes secreted higher levels of LIX, LIF, IL-9, IL-6, GM-CSF, and eotaxin. We note that these signatures cannot be classified as simple delineations of inflammatory or anti-inflammatory cues, but instead reveal complex differences between APOE3 and APOE4 astrocytes in the patterns of interacting immune communications.

**Figure 5.**
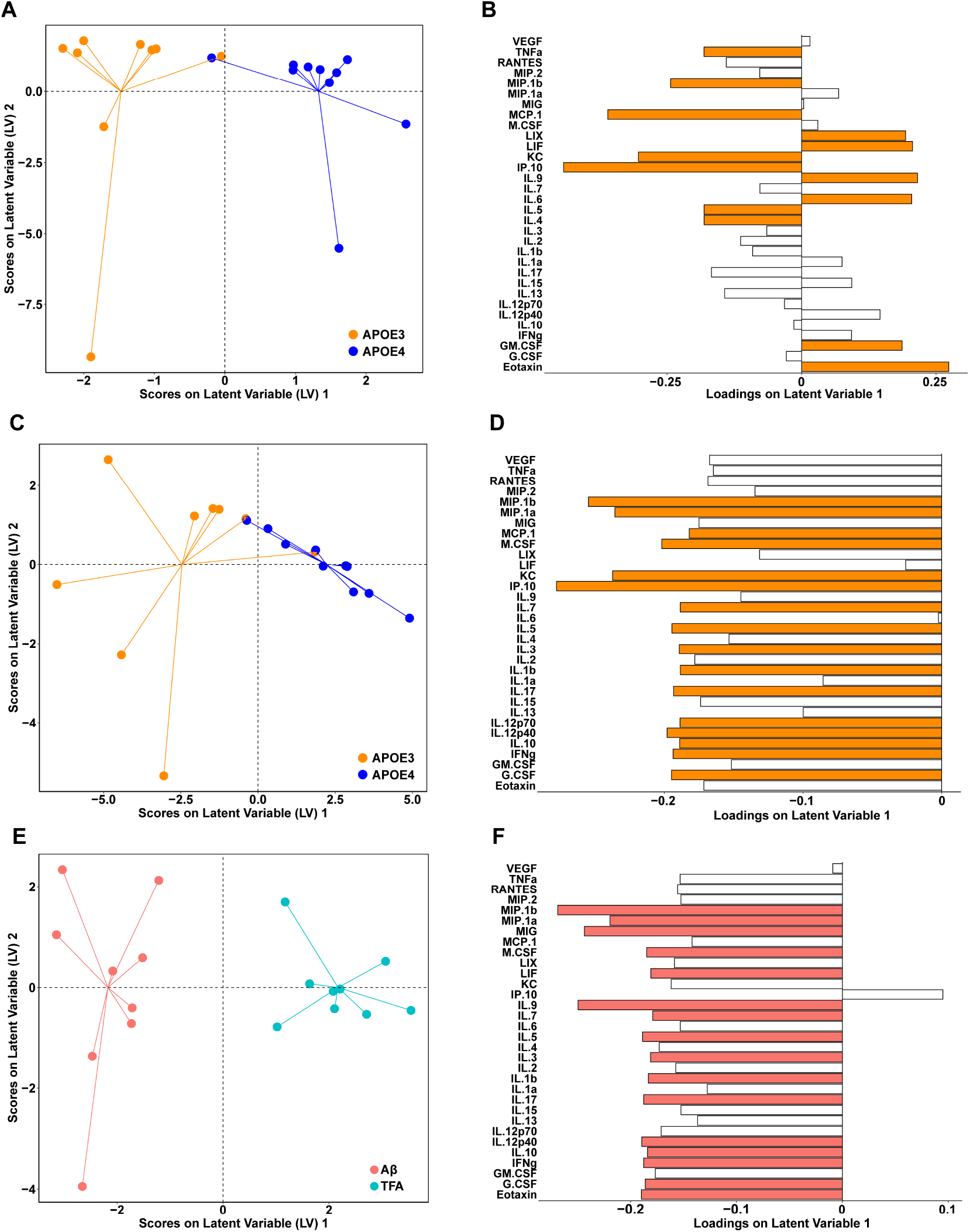
Immune signaling of APOE4 astrocytes lacks response to pathological Aβ. Partial Least Squares Discriminant Analysis (PLSDA) of vehicle treated APOE3 and APOE4 ACM, separated by genotype (Accuracy: 70.17% (5LV); Confidence: 95.06%), scores plot (A) and loadings (B), with VIPs highlighted in orange. PLSDA scores (C) and loadings (D) of 1μM Aβ treated APOE3 and APOE4 ACM, separated by genotype (Accuracy: 72.13% (2LV); Confidence: 98.80%), with VIPs highlighted in orange. PLSDA scores (E) and loadings (F) of APOE3 ACM, separated by treatment with 1 μM Aβ or vehicle (Accuracy: 67.22% (5LV); Confidence: 96.39%), VIPs highlighted in pink.

Having discovered significant differences in cytokine secretion even at rest, we next aimed to define differences in cytokine secretion in response to an AD-relevant stimulus. When we combine all Aβ cytokine reactions for APOE3 and APOE4 astrocytes and predict genotype based on the cytokine secretion pattern of the reactive astrocytes, we observe that, similar to resting state, we are able to distinguish genotype by immune signaling reaction to Aβ stimulation (**Figure 5C and 5D**). With high accuracy (72%) and high confidence (99%), we see that APOE3 astrocytes secrete more of all cytokines measured, consistent with our separate genotype models. VIP (variable importance in projection) cytokines which contribute greater than average to this model, helping separate APOE3 from APOE4 astrocytes treated with Aβ, include MIP-1β, MIP-1α, MCP-1, M-CSF, KC, IP-10, IL-7, IL-5, IL-3, IL-1β, IL-17, IL-12p70 and IL-12p40, IL-10, IFNγ, and G-CSF. These cytokines represent a broad range of functions including inflammation, growth, tissue repair, and neuroprotection.

Focusing on the effects of Aβ stimulation in individual genotypes, we found that APOE3 astrocytes responded robustly to exposure to 1 μM Aβ, increasing levels of nearly all cytokines measured (**Figure 5E and 5F**, model cross-validation accuracy 67%, 96% confidence). Conversely, we could not distinguish the cytokine response of APOE4 astrocytes exposed to Aβ from those treated with vehicle; our PLS model performed no better than random chance (cross-validation accuracy 55%, confidence 54%). Thus, primary APOE3 astrocytes react to Aβ by strongly upregulating general cytokine secretion, while APOE4 astrocytes do not measurably alter the production of immune cues in the presence of Aβ.

Overall, these three models of APOE3 and APOE4 astrocytes at rest and with Aβ stimulation, along with vehicle versus Aβ treatment in APOE3 and APOE4 astrocytes separately, demonstrate that APOE4 astrocytes have an inherent immune signaling divergence from the common APOE3 variant at rest, as well as lack of response to pathological Aβ build up in the extracellular milieu. Since Aβ stimulation did not produce predictable changes in cytokine signaling, a specific cytokine signature is not responsible for the increase in mitochondrial metabolism of APOE4 neurons.

## 4. Discussion

To discover new avenues for mitigating AD risk in APOE4 carriers, we must understand how APOE4 alters responses to AD proteinopathies in a cell-type-specific manner. Astrocytes, the main producers of APOE in the brain, are crucial to supporting neuronal health and resilience, and may hold an important key to identifying mechanisms of neuron vulnerability in AD. Here, we quantified astrocytic live cell glycolytic and oxidative metabolism and immune signaling, two of their main support functions, and found that APOE genotype affects the astrocytic response to Aβ stimulation, which has repercussions affecting neuronal health.

We utilize one-way experiments to isolate the effects of astrocytes on neurons. While astrocyte-neuron co-culture would allow for bi-directional communication between cell types, quantification of the end-state of such cross-communication loses information on directionality of particular signals, leaving us open to misinterpretation of causality and the role of each cell type in driving particular cellular phenomena. Our experimental design enables us to specifically ask whether and how astrocytes are influencing markers of neuronal health.

We found that APOE4 neurons treated with ACM from Aβ-stimulated APOE4 astrocytes reacted with significantly increased basal glycolysis and basal mitochondrial glucose metabolism, while the corresponding APOE3 neurons did not see increases when treated with ACM from Aβ-treated APOE3 astrocytes. Without Aβ stimulation of the astrocytes, addition of ACM from APOE3 and APOE4 astrocytes to corresponding neurons evoked very little increase in metabolic rate. These results differ from those of Qi et al., who found that that stimulating neurons with ACM from corresponding APOE genotype increased maximal respiration of the neurons^27^. While both our study and that of Qi et al. use primary cell cultures derived from the same line of APOE knock-in mice, different length of time in culture (our cells matured longer) or different concentrations of drugs used in Seahorse assays (optimized differently between labs) may have contributed to these discrepancies in outcomes. We followed up on our findings by testing the rate of ATP production in each cell type under each condition and found that while the rate of ATP production in APOE3 neurons was higher than APOE4 neurons in all conditions (data not shown), ATP production in APOE4 astrocytes was higher than in APOE3 astrocytes when treated with Aβ (**Figure 4A**). Astrocyte-generated ATP can activate P2X receptors on post-synaptic neuron terminals to increase AMPA receptor availability, leading to increased neuronal signaling strength^29^. Thus, when we treated APOE4 neurons with media from Aβ-treated APOE4 astrocytes, higher levels of ATP in the media may have led to the rise in neuronal activity, and thus higher APOE4 neuronal metabolic output (**Figure 1C**). This possibility is supported by previous findings by others of a hyperactive phenotype in APOE4 neurons^32^.

Previous studies have quantified the metabolic effects of stimulating primary astrocyte cultures with IL-1β^33^, TNFα^33^, LPS^34^, and Aβ^35^, but these studies did not account for the effect of APOE variants on the downstream metabolic effects of stimulus. APOE is known to modulate Aβ aggregation and clearance in a genotype-dependent manner^36,37^, but to our knowledge, ours is the first study to examine APOE genotype-specific differences in astrocyte metabolic response to Aβ stimulation. Aβ has been found to increase the glucose uptake of primary astrocytes in general, demonstrating increased effects with increasing concentration, as well as with the number of hours of stimulation^35^. Along with increased glucose uptake, production of lactate and glycogen increases when astrocytes are treated with very high concentrations (25 μM) of Aβ for 48 hours^35^. However, we did not observe this increase in astrocytic lactate production with a lower, more disease-physiological concentration of Aβ stimulation. The concentration of Aβ used in cell culture experiments to mimic the environment of the AD brain varies greatly and is often far beyond that found in the human brain, even in end-stage disease^38^. The disease-physiological level of Aβ in brain tissue is cited at 50 nM, with possible localized concentrations of up to 1 μM, however, many studies use up to 50 μM of Aβ to stimulate cells^35^.

Glial activation is a key component of AD^39^ and directly effects cellular glucose and lipid metabolism^40^. We sought to specifically understand the astrocytic APOE genotype differences in cytokine secretion upon Aβ stimulation, since astrocytes are the main producers of APOE in healthy brains and have numerous metabolically supportive functions to serve neurons, including their immune regulation support^12,41–43^. We found that even in the absence of Aβ, APOE4 primary astrocytes secrete a significantly different profile of cytokines compared to their APOE3 counterparts. However, in the presence of Aβ, APOE4 astrocytes did not exhibit a noticeable immune response to the stimulus, unlike the robust increase in general cytokine secretion exhibited by Aβ-treated APOE3 astrocytes. These findings are important because many studies associate APOE4 genotype with increased inflammation^20,37,44^, yet many different measures of inflammation exist, and not all measures are necessarily quantified in every study. For example, tissue swelling, cellular morphological change, and molecular markers like cytokines are all used to describe an inflammatory state, but not all of these measures are necessarily in agreement in a given system. Our finding that APOE4 astrocytes display a lack of change in cytokine secretion profile when stimulated with Aβ does not necessarily transfer to other measures of activation in astrocytes or other APOE4 neural cells and tissue.

In addition to inflammatory processes, cytokine signaling initiates a cascade of downstream intracellular pathways affecting a multitude of cellular processes. Thus, the lack of cytokine secretion response to Aβ we observed in APOE4 astrocytes has profound effects on the ability of cells, including neurons, in the vicinity of those astrocytes to adjust their functions in response to a potential toxic threat. For example, cytokines including TNFα and IL-1β modulate astrocytic glucose utilization, TCA cycle activity, and glycogen levels^45^. Additionally, many cytokine signaling pathways ultimately end in modulating expression in neurons, such as glial TNFα increasing neuronal expression of AMPA receptors^46,47^ which modulate neuron synaptic signaling^47^, and IL-2, IL-3, IL-4, IL-6, IL-10, and IL-12 signaling through JAK/STAT pathways^48,49^ to increase transcription of genes, cell survival, and activation of intracellular pathways such as PI3K/Akt and PKB/Mtor^49^. Thus, modulation of both astrocytic cytokine secretion and metabolism may lead to our observed metabolic changes in neurons.

Recently, single-cell transcriptomic analysis has shined a spotlight on cell-type-specific contributions to AD pathology and progression^50–54^. Astrocytes in particular have been shown to have a ubiquitous role in CNS diseases, which had previously been underappreciated in the field^51^. With their important neuronal support functions, dysfunctional astrocytes contribute to a disease-promoting environment by failing to perform those functions when neurons most need them, such as in the presence of cell stressors like Aβ, leaving neurons more susceptible to injury, dysfunction, and death. Over 500 astrocytic genes are differentially expressed between AD patients and non-demented controls^54^, including genes controlling neuronal synaptic signaling, which were lower in AD astrocytes, and could on a larger scale affect the coordinated firing of neural circuits affecting cognitive function^54^. To our knowledge, no single-cell analysis has been done in brain tissue that identifies differences in astrocytic transcriptomes or proteomes in the context of APOE genotype – a gap waiting to be filled.

Overall, our study demonstrates that APOE4 astrocytes exhibit a lack of cytokine secretion response in the presence Aβ, in tandem with increased glycolytic ATP production as compared to Aβ-exposed APOE3 astrocytes. These differing astrocyte responses to Aβ are carried into the extracellular environment and result in elevated mitochondrial and glycolytic metabolism in APOE4 neurons as compared to APOE3 neurons. This finding may potentially explain increased oxidative stress observed in the brains of human APOE4 carriers^55–57^. Over time, elevated metabolic rate and the consequent increase in oxidative stress and free radical production can result in cellular stress and neuron death. In both healthy aging and in AD, the rates of neuron and synaptic loss increase over time^58^. Thus, increased oxidative stress from APOE4 astrocytes in the presence of Aβ leads to lack of neuronal support and consequent increased risk of neuron synaptic loss, contributing to a disease-promoting environment and increased risk for AD in APOE4 carriers^5,7,8^. These findings in isolated astrocytes and neurons definitively demonstrate astrocyte-specific functional changes in the presence of Aβ, with detrimental downstream effects on neurons. More, these effects vary according to APOE genotype, suggesting mechanisms by which APOE4 creates a disease-promoting environment in the brain by interfering with neuronal support. Our findings highlight the need for investigations of the effects of APOE4 on inter-celltype communications and interactions in the brain to identify new avenues for precision medicine treatment of at-risk APOE4 carriers and AD patients.

## Data Availability Statement

The data of this study are available from the corresponding author upon reasonable request.

## Notes

**Funding Statement**, This work is supported by R01AG072513 from the National Institute on Aging and start-up funds from Penn State College of Medicine Department of Neurosurgery and Department of Pharmacology, both to EAP. RMF is supported by NIH NRSA predoctoral fellowship F31AG071131 from the National Institute on Aging. MKK and DCC are supported by training fellowship T32NS115667 from the National Institute of Neurological Disorders and Stroke.

**Conflict of Interest Disclosure**, The authors declare that they have no conflict of interests with the content of this article.

### Competing Interest Statement

The authors have declared no competing interest.

## References

1. Huang, Y. & Mahley, R. W. Apolipoprotein E: Structure and Function in Lipid Metabolism, Neurobiology, and Alzheimer’s Diseases. Neurobiol Dis 72PA, 3 (2014).

2. Mahley, R. W. Apolipoprotein E: Cholesterol Transport Protein with Expanding Role in Cell Biology. Science (1979) 240, 622–630 (1988).

3. Zannis, V. I., Just, P. W. & Breslow, J. L. Human apolipoprotein E isoprotein subclasses are genetically determined. Am J Hum Genet 33, 11 (1981).

4. Rall, S. C., Weisgraber, K. H. & Mahley, R. W. Human apolipoprotein E. The complete amino acid sequence. Journal of Biological Chemistry 257, 4171–4178 (1982).

5. Farrer, L. A. et al. Effects of Age, Sex, and Ethnicity on the Association Between Apolipoprotein E Genotype and Alzheimer Disease: A Meta-analysis. JAMA 278, 1349–1356 (1997).

6. Belloy, M. E., Napolioni, V. & Greicius, M. D. A Quarter Century of APOE and Alzheimer’s Disease: Progress to Date and the Path Forward. Neuron 101, 820–838 (2019).

7. Corder, E. H. et al. Gene dose of apolipoprotein E type 4 allele and the risk of Alzheimer’s disease in late onset families. Science 261, 921–3 (1993).

8. Strittmatter, W. J. & Roses, A. D. Apolipoprotein E and Alzheimer’s Disease. Annu Rev Neurosci 19, 53–77 (1996).

9. Mahley, R. W. Central Nervous System Lipoproteins. Arterioscler Thromb Vasc Biol 36, 1305–1315 (2016).

10. Perkins, M. et al. Altered Energy Metabolism Pathways in the Posterior Cingulate in Young Adult Apolipoprotein E ε4 Carriers. J Alzheimers Dis 53, 95–106 (2016).

11. Verkhratsky, A., Nedergaard, M. & Hertz, L. Why are Astrocytes Important? Neurochem Res 40, 389–401 (2015).

12. Prebil, M., Jensen, J., Zorec, R. & Kreft, M. Astrocytes and energy metabolism. Arch Physiol Biochem 117, 64–69 (2011).

13. Choi, S. S., Lee, H. J., Lim, I., Satoh, J. I. & Kim, S. U. Human astrocytes: Secretome profiles of cytokines and chemokines. PLoS One 9, (2014).

14. Verkhratsky, A., Olabarria, M., Noristani, H. N., Yeh, C. Y. & Rodriguez, J. J. Astrocytes in Alzheimer’s disease. Neurotherapeutics 7, 399–412 (2010).

15. Boyles, J. K., Pitas, R. E., Wilson, E., Mahley, R. W. & Taylor, J. M. Apolipoprotein E associated with astrocytic glia of the central nervous system and with nonmyelinating glia of the peripheral nervous system. J Clin Invest 76, 1501–1513 (1985).

16. de Chaves, E. P. & Narayanaswami, V. Apolipoprotein E and cholesterol in aging and disease in the brain. Future Lipidol 3, 505–530 (2008).

17. Ioannou, M. S. et al. Neuron-Astrocyte Metabolic Coupling Protects against Activity-Induced Fatty Acid Toxicity. Cell 177, 1522–1535.e14 (2019).

18. Sienski, G. et al. APOE4 disrupts intracellular lipid homeostasis in human iPSC-derived glia. Sci Transl Med 13, (2021).

19. Lanfranco, M. F., Sepulveda, J., Kopetsky, G. & Rebeck, G. W. Expression and secretion of apoE isoforms in astrocytes and microglia during inflammation. Glia 69, 1478–1493 (2021).

20. Leeuw, S. M. de et al. APOE2, E3, and E4 differentially modulate cellular homeostasis, cholesterol metabolism, and inflammatory response in isogenic iPSC-derived astrocytes. Stem Cell Reports 0, (2021).

21. Liu, C. C., Kanekiyo, T., Xu, H. & Bu, G. Apolipoprotein e and Alzheimer disease: Risk, mechanisms and therapy. Nat Rev Neurol 9, 106–118 (2013).

22. Bennett, D. A. et al. Neuropathology of older persons without cognitive impairment from two community-based studies. Neurology 66, 1837–1844 (2006).

23. Nicholas, D. et al. Advances in the quantification of mitochondrial function in primary human immune cells through extracellular flux analysis. PLoS One 12, e0170975 (2017).

24. Thévenot, E. A., Roux, A., Xu, Y., Ezan, E. & Junot, C. Analysis of the Human Adult Urinary Metabolome Variations with Age, Body Mass Index, and Gender by Implementing a Comprehensive Workflow for Univariate and OPLS Statistical Analyses. J Proteome Res 14, 3322–3335 (2015).

25. Villanueva, R. A. M. & Chen, Z. J. ggplot2: Elegant Graphics for Data Analysis (2nd ed.). Measurement (Mahwah N J) 17, 160–167 (2019).

26. Wu, L., Zhang, X. & Zhao, L. Human ApoE Isoforms Differentially Modulate Brain Glucose and Ketone Body Metabolism: Implications for Alzheimer’s Disease Risk Reduction and Early Intervention. The Journal of Neuroscience 38, 6665–6681 (2018).

27. Qi, G. et al. ApoE4 Impairs Neuron-Astrocyte Coupling of Fatty Acid Metabolism. Cell Rep 34, (2021).

28. Koizumi, S., Fujishita, K. & Inoue, K. Regulation of cell-to-cell communication mediated by astrocytic ATP in the CNS. Purinergic Signal 1, 211 (2005).

29. Boué-Grabot, E. & Pankratov, Y. Modulation of Central Synapses by Astrocyte-Released ATP and Postsynaptic P2X Receptors. Neural Plast 2017, (2017).

30. Gavillet, M., Allaman, I. & Magistretti, P. J. Modulation of astrocytic metabolic phenotype by proinflammatory cytokines. Glia 56, 975–989 (2008).

31. Martens, Y. A. et al. ApoE Cascade Hypothesis in the pathogenesis of Alzheimer’s disease and related dementias. Neuron (2022) doi:10.1016/J.NEURON.2022.03.004.

32. Nuriel, T. et al. Neuronal hyperactivity due to loss of inhibitory tone in APOE4 mice lacking Alzheimer’s disease-like pathology. Nature Communications 2017 8:1 8, 1–14 (2017).

33. Jiang, T. & Cadenas, E. Astrocytic metabolic and inflammatory changes as a function of age. Aging Cell 13, 1059–1067 (2014).

34. Robb, J. L. et al. The metabolic response to inflammation in astrocytes is regulated by nuclear factor-kappa B signaling. Glia 68, 2246–2263 (2020).

35. Allaman, I. et al. Amyloid-β Aggregates Cause Alterations of Astrocytic Metabolic Phenotype: Impact on Neuronal Viability. Journal of Neuroscience 30, 3326–3338 (2010).

36. Holtzman, D. M. et al. Apolipoprotein E isoform-dependent amyloid deposition and neuritic degeneration in a mouse model of Alzheimer’s disease. Proc Natl Acad Sci U S A 97, 2892–2897 (2000).

37. Liu, C.-C. et al. ApoE4 Accelerates Early Seeding of Amyloid Pathology. Neuron 96, 1024–1032.e3 (2017).

38. LaRocca, T. J. et al. Amyloid beta acts synergistically as a pro-inflammatory cytokine. Neurobiol Dis 159, 105493 (2021).

39. Heneka, M. T. et al. Neuroinflammation in Alzheimer’s disease. Lancet Neurol 14, 388–405 (2015).

40. Shi, J., Fan, J., Su, Q. & Yang, Z. Cytokines and Abnormal Glucose and Lipid Metabolism. Front Endocrinol (Lausanne) 10, 703 (2019).

41. Sofroniew, M. v. & Vinters, H. v. Astrocytes: biology and pathology. Acta Neuropathol 119, 7–35 (2010).

42. Pellerin, L. & Magistretti, P. J. Glutamate uptake into astrocytes stimulates aerobic glycolysis: a mechanism coupling neuronal activity to glucose utilization. Proceedings of the National Academy of Sciences 91, 10625–10629 (1994).

43. Magistretti, P. J. & Pellerin, L. Astrocytes couple synaptic activity to glucose utilization in the brain. News in Physiological Sciences 14, 177–182 (1999).

44. Shi, Y. et al. ApoE4 markedly exacerbates tau-mediated neurodegeneration in a mouse model of tauopathy. Nature 549, 523–527 (2017).

45. Gavillet, M., Allaman, I. & Magistretti, P. J. Modulation of astrocytic metabolic phenotype by proinflammatory cytokines. Glia 56, 975–989 (2008).

46. Beattie, E. C. et al. Control of Synaptic Strength by Glial TNFα. Science (1979) 295, 2282–2285 (2002).

47. Stellwagen, D. & Malenka, R. C. Synaptic scaling mediated by glial TNF-α. Nature 2006 440:7087 440, 1054–1059 (2006).

48. Morris, R., Kershaw, N. J. & Babon, J. J. The molecular details of cytokine signaling via the JAK/STAT pathway. Protein Sci 27, 1984 (2018).

49. Gadani, S. P., Cronk, J. C., Norris, G. T. & Kipnis, J. IL-4 in the Brain: A Cytokine To Remember. The Journal of Immunology 189, 4213–4219 (2012).

50. Mathys, H. et al. Single-cell transcriptomic analysis of Alzheimer’s disease. Nature 570, 332–337 (2019).

51. Batiuk, M. Y. et al. Identification of region-specific astrocyte subtypes at single cell resolution. Nature Communications 2020 11:1 11, 1–15 (2020).

52. Bayraktar, O. A. et al. Astrocyte layers in the mammalian cerebral cortex revealed by a single-cell in situ transcriptomic map. Nature Neuroscience 2020 23:4 23, 500–509 (2020).

53. Hasel, P., Rose, I. V. L., Sadick, J. S., Kim, R. D. & Liddelow, S. A. Neuroinflammatory astrocyte subtypes in the mouse brain. Nature Neuroscience 2021 24:10 24, 1475–1487 (2021).

54. Lau, S. F., Cao, H., Fu, A. K. Y. & Ip, N. Y. Single-nucleus transcriptome analysis reveals dysregulation of angiogenic endothelial cells and neuroprotective glia in Alzheimer’s disease. Proc Natl Acad Sci U S A 117, 25800–25809 (2020).

55. Butterfield, D. A. & Mattson, M. P. Apolipoprotein E and oxidative stress in brain with relevance to Alzheimer’s disease. Neurobiol Dis 138, 104795 (2020).

56. Tönnies, E. & Trushina, E. Oxidative Stress, Synaptic Dysfunction, and Alzheimer’s Disease. Journal of Alzheimer’s Disease 57, 1105 (2017).

57. Cioffi, F., Adam, R. H. I. & Broersen, K. Molecular Mechanisms and Genetics of Oxidative Stress in Alzheimer’s Disease. J Alzheimers Dis 72, 981–1017 (2019).

58. Long, J. M. & Holtzman, D. M. Alzheimer Disease: An Update on Pathobiology and Treatment Strategies. Cell 179, 312–339 (2019).

